# Feedback regulation of iron-sulfur cluster biogenesis

**DOI:** 10.1101/2025.06.15.659787

**Authors:** Stephanie M. Stuteley, Jiahua Chen, Jin Wang, Stephanie Dawes, Edward N. Baker, Christopher J. Squire, Maria-Eirini Pandelia, Ghader Bashiri

## Abstract

Iron-sulfur (Fe-S) clusters are ubiquitous cofactors in biological systems. Given their central role in bacterial metabolism and pathogenesis, the biogenesis of Fe-S clusters is tightly controlled. We reveal a feedback regulatory mechanism involving the sulfide producing SufS/SufU complex within the sulfur utilization (SUF) system of *Mycobacterium tuberculosis*, the bacterium that causes tuberculosis. In this mechanism, [2Fe-2S] clusters compete with zinc ions for binding to the sulfide transfer protein SufU. Cluster binding induces SufU tetramerization, which prevents its interaction with the cysteine desulfurase SufS, thereby inhibiting SufS activation and limiting sulfide supply for Fe-S cluster biogenesis. These findings uncover an unrecognized regulatory mechanism in *M. tuberculosis*, ensuring strict control of Fe-S cluster production.

## Introduction

Iron-sulfur (Fe-S) clusters are ubiquitous and highly versatile protein cofactors present in all domains of life.^1^ These clusters are utilized by Fe-S proteins in many essential biological processes, such as redox sensing, catalysis, and electron transfer.^2^ Fe-S cluster formation involves a complex network of proteins that mobilize sulfur and iron, assemble nascent clusters, and transfer them to target Fe-S cluster proteins. In bacteria, this process is orchestrated by sophisticated Fe-S biogenesis systems, including the sulfur utilization (SUF), iron-sulfur cluster (ISC), and nitrogen fixation (NIF) systems.^3^ Of these, the SUF system displays the broadest distribution in bacteria^4^ and is found in *Mycobacterium tuberculosis*,^5^ the causative agent of tuberculosis. Notably, the SUF system in *M. tuberculosis* is implicated in pathogenesis, virulence, and the transition to a dormant, non-replicating state.^6–8^

The SUF system in *M. tuberculosis* is encoded by the *sufRBDCSUT* operon, comprising *rv1460-1466* open reading frames (ORFs), and is co-transcribed as a single mRNA transcript.^5^ Fe-S cluster biogenesis begins with the joint action of the cysteine desulfurase SufS (Rv1464) and the sulfide transfer protein SufU (Rv1465), which provide sulfur for cluster formation.^9,10^ SufU forms a zinc-mediated complex with SufS, enabling the transfer of sulfide from SufS to SufU.^9,11^ This interaction activates SufS by efficiently removing the abstracted sulfide, which is thought to be the rate-limiting step in the turnover of active sites in *E. coli* SufS.^12^ SufU then delivers the extracted sulfide to the SufB component of the SufBCD core scaffold complex (Rv1461-Rv1463), where it is proposed that cluster assembly occurs.^10,13,14^ This direct transfer mechanism avoids the release of toxic sulfide during Fe-S cluster assembly.

The extreme susceptibility of Fe-S clusters to oxidation, along with the inherent toxicity associated with free iron and sulfur, necessitates strict regulation of cluster assembly to avoid futile cycling and metabolic overload. In *M. tuberculosis*, the SUF machinery is controlled through both transcriptional and post-translational mechanisms. SufR (Rv1460) acts as a transcriptional repressor that regulates the expression of the suf operon by binding [4Fe-4S] clusters.^15,16^ In addition, SufB (Rv1461) contains a unique intein, a self-splicing protein insert found exclusively in pathogenic mycobacteria.^17^ The intein functions as a sensor for oxidative and nitrosative stress in *M. tuberculosis*,^18^ and its post-translational splicing is essential for assembling the core SufBCD complex.^5,17^

Here, we describe an inhibitory feedback mechanism within the SUF system, driven by the alternate binding of zinc ions or Fe-S clusters to SufU. Specifically, zinc binding facilitates the formation the SufS/SufU complex, while Fe-S cluster binding inhibits this interaction and prevents SufS activation, thereby restricting the sulfur supply for Fe-S biogenesis. This third regulatory mechanism highlights a multi-layered regulatory system that ensures Fe-S cluster production is tightly modulated in response to environmental signals and cellular demands.

## Results

### *Mtb*-SufU binds [2Fe-2S]^2+^ clusters

Over-expression of *Mtb*-SufU in *E. coli* BL21 (DE3) cells produced two distinct SufU species. The anticipated monomeric SufU protein (17.7 kDa) bound to zinc^9,11,19^ was confirmed by size-exclusion chromatography coupled with multi-angle laser light scattering (SEC-MALS) (Fig. 1A, blue trace) and inductively-coupled plasma mass spectrometry (ICP-MS) analysis (Table S1). Unexpectedly, a tetrameric form of SufU species was also present in the SEC chromatograms and confirmed by SEC-MALS (Fig. 1A, pink trace). The UV/Vis spectrum of the SufU tetramer displayed features typical of [2Fe-2S] clusters (Fig. 1B).

**Fig. 1.**
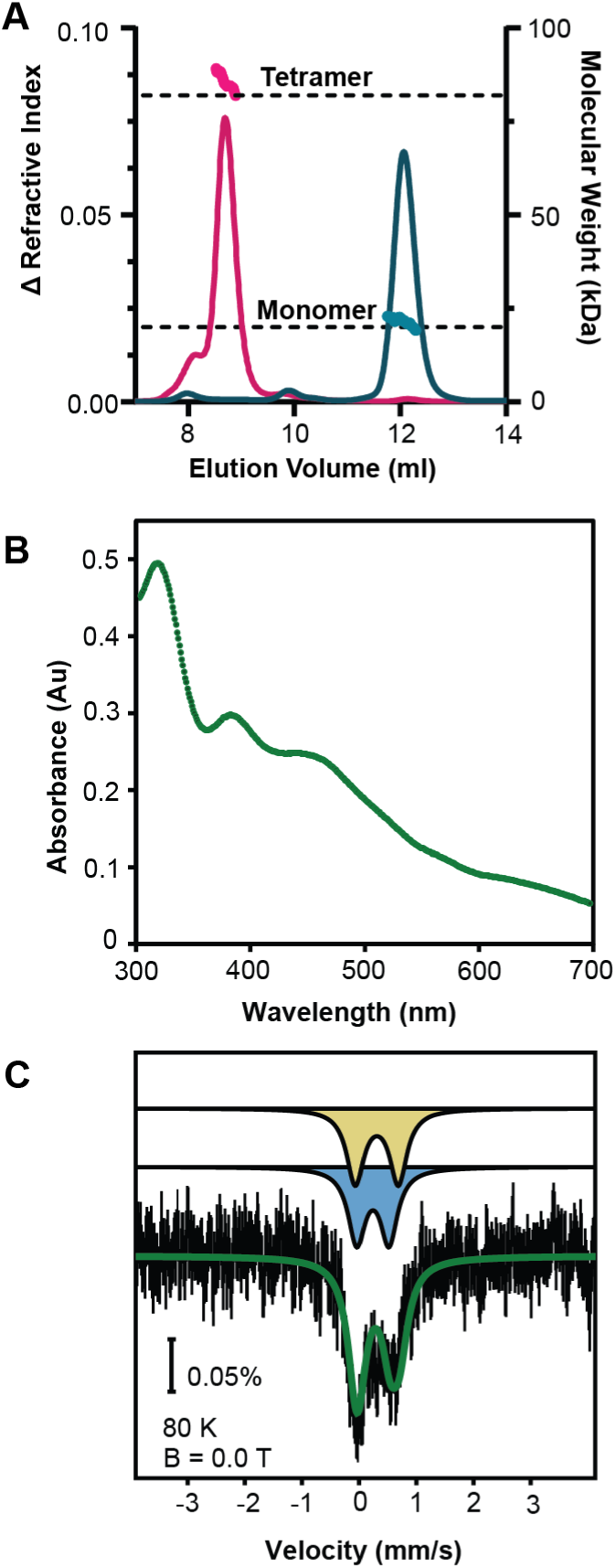
*Mtb*-SufU binds [2Fe-2S]^2+^ clusters. **(A)** SEC MALS analysis of [2Fe-2S]-SufU suggests tetramer formation (magenta), whereas Zn-SufU exists as a monomer in solution (teal). The refractive index for each sample is shown as solid colored lines, with the average molecular weight of the samples shown as continuous dots. **(B)** UV/Vis spectrum of as-purified *Mtb*-SufU, showing characteristic peaks of a [2Fe-2S] cluster with absorbances at 320 nm, 385 nm, and 460 nm. **(C)** Mössbauer spectrum of as-purified *Mtb*-SufU was recorded at 80 K in the absence of an external magnetic field. The total simulation, shown as a green line, represents the sum of subspectra from two Fe³⁺ ions in a [2Fe-2S] cluster, depicted in yellow (δ = 0.24 mm/s, ΔEQ = 0.56 mm/s) and blue (δ = 0.31 mm/s, ΔEQ = 0.75 mm/s).

While the cluster-bound SufU constituted ∼10% of the *Mtb*-SufU produced in *E. coli* cells, this ratio could be shifted to ∼75% of the total SufU when expressed using iron-enriched protocols, including co-expression with the *E. coli* suf operon^20^ and in media supplemented with iron and sulfur sources.

We next used ^57^Fe Mössbauer spectroscopy to validate the presence of [2Fe-2S] clusters. Since [2Fe-2S]-SufU could not be produced in minimal media, the protein was expressed in Terrific Broth media and Mössbauer spectroscopy was carried out on anaerobically purified protein samples, taking advantage of the 2.1% natural abundance of ^57^Fe. This method excludes [4Fe-4S] cluster formation that can occur during *in vitro* chemical reconstitution, based on the well-documented behavior of cluster assembly in structurally homologous proteins.^21,22^ The Mössbauer spectrum (Fig. 1C), while weak in intensity, is best fit by a set of quadrupole doublets of equal integrated intensity with the following parameters: δ_1_ = 0.24 mm/s, ΔE_Q1_ = 0.56 mm/s and δ_2_ = 0.31 mm/s, ΔE_Q2_ = 0.75 mm/s. These values are characteristic of [2Fe-2S] clusters with two distinct Fe³⁺ sites: the first coordinated by thiol groups and the second bound to a non-thiol ligand. The observed parameters align with those reported for the [2Fe-2S]²⁺ cluster in *E. coli* IscU (*Ec*-IscU),^23^ a known iron-sulfur scaffold protein that shares structural homology with *Mtb*-SufU.

### [2Fe-2S]-bound SufU modulates SufS activity

Next, we performed cysteine desulfurase activity assays to determine the effect of [2Fe-2S]-SufU on SufS activation. To this end, we developed a gas chromatography-mass spectrometry (GC-MS) method to directly measure L-Ala production as an indicator of L-Cys consumption by SufS. This GC-MS assay eliminates interference from Fe-S clusters in the classical sulfide release assay, which indirectly monitors sulfide release from L-Cys via methylene blue formation. In these anaerobic experiments, SufS was pre-incubated with either Zn^2+-^bound or [2Fe-2S]-bound SufU prior to L-Cys addition to initiate the reaction.

While the addition of Zn^2+-^SufU increased the rate of L-Cys conversion to L-Ala by 10-fold, the reaction with [2Fe-2S]-SufU showed a rate comparable to the SufS only reaction (Fig 2). These results demonstrate that the Fe-S cluster adversely impacts SufS activation, likely by obstructing the interaction between SufU and SufS.

**Fig. 2.**
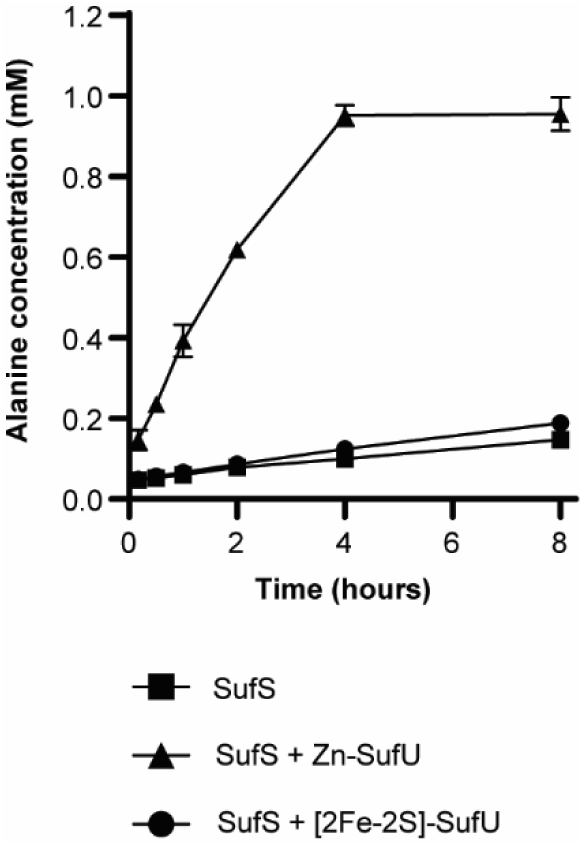
[2Fe-2S] clusters abolish SufU interaction with SufS. A GC-MS functional assay monitors the production of L-Ala as a product of the cysteine desulfurase activity of SufS. *Mtb*-SufS shows low basal activity that increases 10-fold in the presence of two molar equivalents of Zn-bound *Mtb*-SufU after 4 hours. In contrast, [2Fe-2S]-bound *Mtb*-SufU does not activate the cysteine desulfurase activity of SufS under the same conditions. Error bars indicate standard deviation in triplicates.

To establish the molecular mechanism underlying this effect, we determined the crystal structures of a [2Fe-2S]-bound *Mtb*-SufU and a Zn^2+-^mediated *Mtb*-SufS/SufU complex at 1.95 and 2.0 Å resolution, respectively (Fig. 3A, Fig. S1, Table S2). While clear and unambiguous electron density revealed binding of a [2Fe-2S] cluster in the metal binding pocket of SufU (Fig. 3B), the overall [2Fe-2S]-SufU structure remained highly similar to the Zn^2+^-SufU seen in the SufS/SufU complex (0.62 Å root mean square deviation, RMSD, over 135 Cα positions) (Fig. 3C). In the SufS/SufU structure, the Asp42, Cys67, and Cys131 residues from SufU, and His354 from SufS, coordinate Zn^2+^, bridging the two proteins (Fig 3C). This Zn^2+^-mediated interaction facilitates the sulfide transfer from SufS-Cys373 to SufU-Cys40, similarly to SufS/SufU in *B. subtilis* (SufS-Cys361 and SufU-Cys41) and *S. aureus* (SufS-Cys389 and SufU-Cys43). ^9,11,19^ The [2Fe-2S]-bound *Mtb*-SufU structure revealed that the [2Fe-2S] cluster is also coordinated by Asp42, Cys67, and Cys131, with SufU-Cys40 acting as the fourth ligand (Fig. 3C), consistent with the Mössbauer spectroscopy results on adopting one non-thiol coordination (Fig. 1C). Together, these observations demonstrate that the [2Fe-2S]^2+^ cluster occupies the same binding site as Zn^2+^ in SufU.

**Fig. 3.**
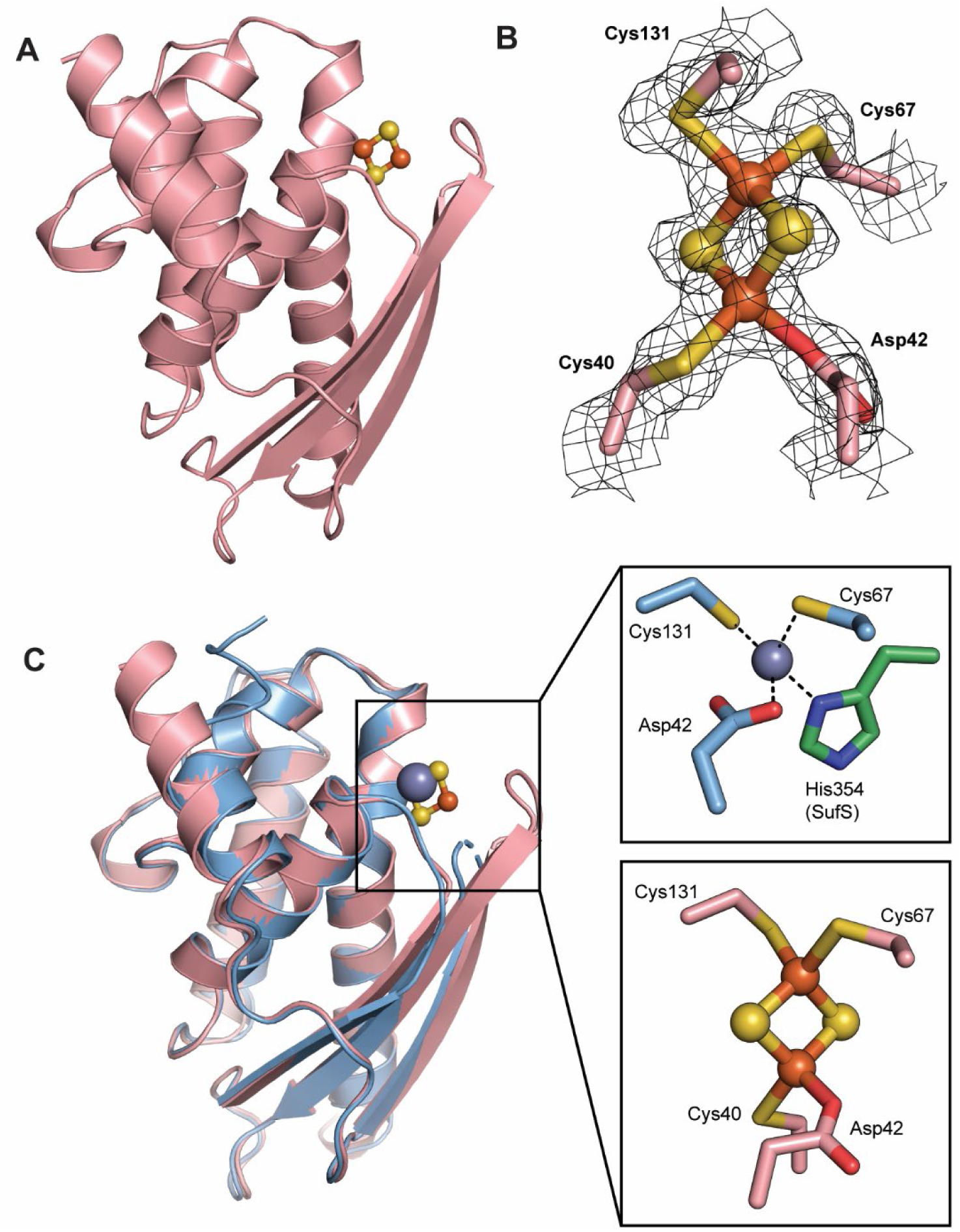
Crystal structure of *Mtb*-SufU. **(A)** SufU binds a [2Fe-2S] cluster in the Zn binding pocket. **(B)** Unbiased 2*Fo*-*Fc* electron density map contoured at σ=1.0, showing coordination of [2Fe-2S] cluster. **(C)** Crystal structure overlay of [2Fe-2S]-SufU (pink) with the Zn^2+^- SufU (blue) from the SufSU complex structure reveals a highly similar overall structure, with a root mean square deviation (RMSD) of 0.62 Å over 135 Cα positions. The insets show the coordination of either a Zn^2+^ ion (gray sphere) or a [2Fe-2S] cluster. In the [2Fe-2S]-SufU structure, SufS-His354 is substituted by SufU-Cys40.

The transition from monomeric Zn^2+-^bound SufU to tetrameric [2Fe-2S]-bound SufU disrupts its interaction with SufS. This tetrameric structure (Fig. 4) is stabilized by multiple polar interactions involving the cluster-forming residues Cys40, Asp42, and Cys67, along with adjacent residues Ile39, Ser68, and Arg128. These residues interact with the open face of a β-strand from another monomer, comprising Phe29, Glu27, Gln32, and Tyr34 (Fig. 4 inset). This oligomeric arrangement creates a solvent-secluded pocket between neighboring monomers, effectively encapsulating the [2Fe-2S] clusters. This encapsulation restricts access to the clusters and prevents any interaction with SufS. The ability of [2Fe-2S] clusters to drive SufU oligomerization represents a unique regulatory mechanism, distinct from their role in homologous IscU-type proteins.^24,25^

**Fig. 4.**
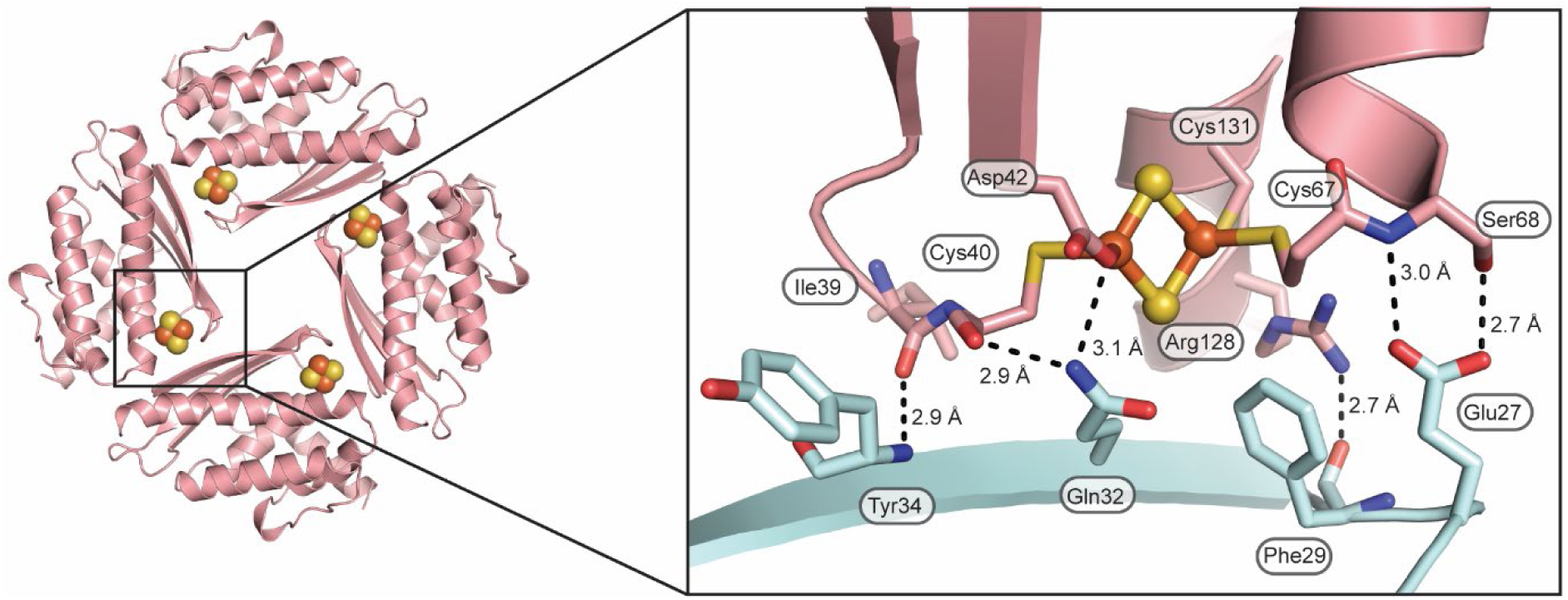
[2Fe-2S]-SufU tetramer structure and interface. The [2Fe-2S]-SufU tetramer structure is generated by applying crystallographic symmetry and is predicted by the PDBePISA server.^47^ The inset highlights polar interactions between adjacent monomers, colored in pink and cyan. The three residues coordinating the clusters, Cys40, Asp42, and Cys67, and adjacent residues contribute to the formation of the tetrameric structure, shielding the [2Fe-2S] clusters within a pocket formed between neighboring monomers.

### Feedback regulation of the SUF system

Our data highlight a novel function for *Mtb*-SufU, beyond its roles in enhancing the low basal cysteine desulfurase activity of SufS and shuttling the acquired sulfides to the core complex for Fe-S cluster biogenesis. To determine if this new function of SufU is conserved in other bacteria, we investigated SufU homologs from *B. subtilis* (*Bs*-SufU) and *S. aureus* (*Sa*-SufU). *Bs*-SufU and *Sa*-SufU share 33.5% and 36.5% sequence identity with *Mtb*-SufU, respectively, with conserved residues involved in metal coordination. Similarly to *Mtb*-SufU, *Bs*-SufU is required for SufS activation,^11,26,27^ while *Sa*-SufU only increases SufS activity by 1.5-fold.^19^ Over-expression of *Sa*-SufU in *E. coli* host cells using iron-enriched protocols produced a colorless protein exclusively, loaded with 1.1 molar equivalents of zinc as confirmed by ICP-MS (Table S1). Conversely, under the same conditions, *Bs*-SufU was purified with either zinc or iron, as observed by ICP-MS (Table S1). However, these results showed only ∼0.5 Fe molar equivalents per SufU monomer (Table S1). While we attempted to chemically reconstitute *Bs*-SufU, it does not appear that the protein can stably bind to the clusters as inferred from the lack of characteristic UV/Vis features. These findings agree with previous *in vitro Bs*-SufU reconstitution experiments, suggesting these results were non- physiological ^28^ or that the *E. coli* host is unable to interact with and load a stable cluster onto *Bs*-SufU. The iron-enriched *Bs*-SufU, similarly to the [2Fe-2S]-cluster *Mtb*-SufU, could not activate *Bs*-SufS (Fig. S2B).

The role of SufU proteins in binding Fe-S clusters and their potential function as scaffold proteins have been controversial in the literature. This arises from comparisons with the well-studied ISC system in *E. coli*, where *Ec-*IscU acts as the scaffold for Fe-S cluster assembly. However, IscU does not appear functionally equivalent to SufU proteins. Genetic complementation studies support this, showing that *E. coli iscSU* cannot restore the *B. subtilis* Δ*sufSU* mutant phenotype.^29^ We were unable to form Fe-S clusters on *Mtb*-SufU *in vitro* using purified Zn^2+^-SufU and SufS. We were able to chemically reconstitute [4Fe-4S] clusters on [2Fe-2S]-*Mtb*-SufU, however, these reverted over time to [2Fe-2S] clusters, as indicated by UV/Vis absorption spectra. Based on these findings, we conclude that *Mtb*-SufU is a bona fide Fe-S binding protein but does not serve as a scaffold for Fe-S cluster formation.

## Conclusions

We propose a feedback regulation mechanism for Fe-S cluster biogenesis in the SUF system (Fig. 5). Under conditions of excess Fe-S clusters, [2Fe-2S] clusters occupy the zinc binding site in SufU, blocking the SufS/SufU interaction and preventing activation of SufS. This leads to reduced sulfide production by SufS, thereby limiting Fe-S cluster biogenesis. This proposed regulatory mechanism operates alongside the transcriptional regulator SufR, which represses transcription of the suf operon when bound to [4Fe-4S] clusters, further reducing Fe-S cluster formation under similar conditions. SufU and SufR bind [2Fe-2S] and [4Fe-4S] clusters, respectively, suggesting they may respond differently to environmental cues. Indeed, we demonstrated that *Mtb*-SufR and *Mtb*-SufU exhibit distinct responses to oxidative stress induced by H_2_O_2_ treatment (Fig. S3). Although [2Fe-2S] clusters are generally more resistant to oxidation than [4Fe-4S] clusters,^30^ the formation of the [2Fe-2S]- SufU tetramer, which encases and shields the [2Fe-2S] clusters, provides additional protection. This protective feature likely facilitated the crystallization of [2Fe-2S]-SufU under aerobic conditions in our study.

**Fig. 5.**
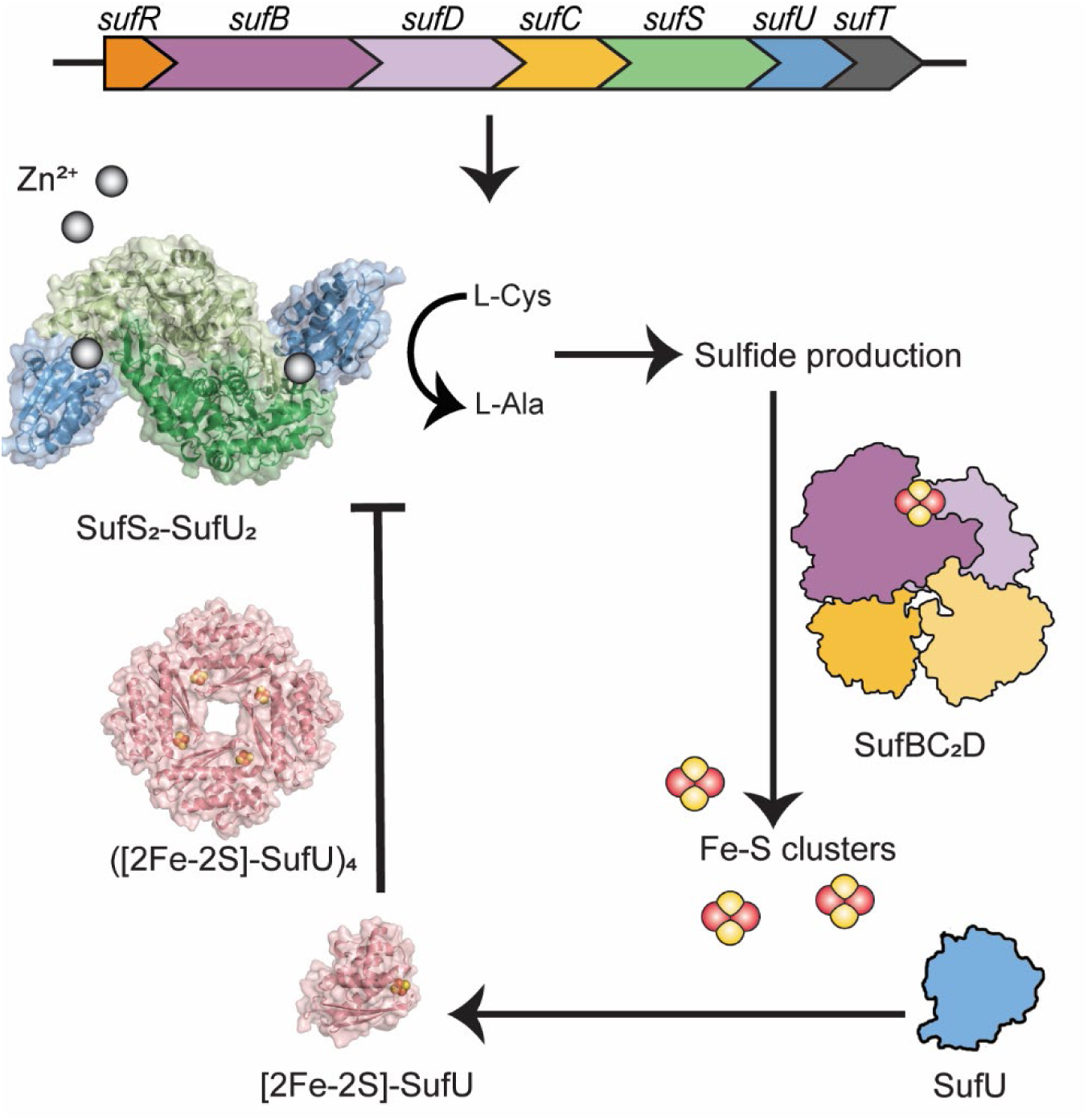
Schematic representation of the proposed feedback regulation mechanism in the SUF machinery of *M. tuberculosis*. Under conditions of cellular demand for Fe-S clusters, SufU binds to zinc, which facilitates it interaction with SufS. This interaction activates the cysteine desulfurase activity of SufS, supplying sulfide for Fe-S cluster assembly on the SufBCD core complex. Once sufficient clusters are formed, excess clusters bind to SufU, prompting the formation of a tetramer. This tetrameric state prevents SufU from interacting with SufS, thereby inhibiting further sulfide production for Fe-S cluster formation.

Under stress conditions such as oxidative/nitrosative stress or iron starvation, disruption of the SufR clusters leads to de-repression of the suf operon, which enhances Fe-S cluster production. It remains unclear, however, if these stress conditions also result in [2Fe- 2S] cluster removal from the existing pool of SufU proteins. Nonetheless, the subsequent production of SufU following SufR de-repression could lead to the formation of Zn^2+^-SufU, which interacts with SufS and enhances Fe-S cluster production. Once the stressors are removed or sufficient clusters are formed, SufR can bind [4Fe-4S] clusters and repress transcription, while cluster loaded SufU prevents its interaction with SufS. Together, these processes ultimately slow down cluster production.

It remains elusive why *M. tuberculosis* evolved SufU to introduce an additional layer of complexity to Fe-S cluster biogenesis. One plausible explanation, beyond its regulatory role, is that SufU binding to [2Fe-2S] clusters acts as a mechanism to sequester excess Fe-S clusters under fluctuating environmental conditions, thereby protecting the cells from potential damage. This adaptation may reflect the pathogen’s ability to survive hypoxic conditions within the host cells by preserving the clusters until transcription regulation takes effect. This is particularly intriguing given that *M. tuberculosis* relies heavily on Fe-S cluster proteins for its metabolism, with the frequency of Fe-S cluster motifs (6.5 motifs/1000 open reading frames) more closely resembling anaerobic bacteria (7.4 motifs) than aerobic (2.8 motifs) or facultative anaerobic (3.7 motifs) bacteria.^7,31^ Our findings highlight the critical role of Fe-S cluster regulation in *M. tuberculosis*, employing multiple layers of control mechanisms to adapt to its niche host environment.

## Materials and Methods

### Protein Expression and Purification

#### SufU constructs

The ORF encoding SufU of *M. tuberculosis* H37Rv was amplified from genomic DNA, then cloned into a pProEX-HTb vector. The ORFs encoding SufU proteins from *B. subtilis* and *S. aureus* were synthesized and cloned (GenScript) into pProEx HTb vectors, then expressed in *E. coli* LOBSTR BL21 (DE3) cells^32^ alone or co-expressed with the *E. coli sufABCDSE* operon.^20^ The N-terminal His_6_-tagged protein was purified by immobilized metal affinity chromatography (IMAC) using Talon resin (Clontech, Takara Bio.) and size-exclusion chromatography (SEC) on a Superdex 75 10/30 column (Cytiva), in a base buffer of 20 mM HEPES (pH 7.5), 200 mM NaCl, 5% glycerol, and 5 mM β-mercaptoethanol (β-Me). Proteins were purified within an mBraun anaerobic glovebox, maintained at <0.1 ppm oxygen concentration.

#### SufS constructs

The ORFs encoding SufS proteins from *M. tuberculosis* H37Rv, *B. subtilis*, and *S. aureus* were synthesized and cloned (GenScript) into the pYUB-Ex vector.^33^ These constructs express His_6_-tagged proteins, which were over-expressed in *M. smegmatis* mc^2^4517 cells.^34^ The cells were grown in a fermenter (BioFlo®415, New Brunswick Scientific) for 4 days and proteins purified as described for SufU, with the SEC step carried out using a Superdex 200 10/30 column (Cytiva).

#### Mtb-SufR

The ORF encoding SufR of *M. tuberculosis* H37Rv was amplified from the genomic DNA and cloned into the pProEx-HTb plasmid which encodes an N-terminal His_6_-tag. Expression was carried out in *E. coli* LOBSTR BL21 (DE3) cells^32^ with co-expression with the *E. coli sufABCDSE* operon.^20^ The cell pellets were mechanically lysed in a buffer containing 50 mM HEPES (pH 7.5), 300 mM KCl, 150 mM NaCl, 4 mM imidazole, 10% glycerol, and 14 mM β-Me, before loading onto a HiTrap Ni-NTA column. After washing to remove non-specific proteins, an Fe-S cluster reconstitution step was performed on the immobilized protein.^35,36^ Five column volumes of reconstitution buffer (0.4 mM Na_2_S, 0.6 mM FeCl_3_, 50 mM HEPES (pH 7.5), 300 mM KCl, 10 % glycerol, 50 mM β-Me) were passed over the column, followed by washing with the lysis buffer. Reconstituted SufR was then eluted from the affinity column and further purified by SEC as above using a buffer containing 20 mM HEPES (pH 8.0), 300 mM KCl, 10 % glycerol, and 5 mM 1,4-dithiothreitol (DTT). Protein was purified within an mBraun anaerobic glovebox, maintained at <0.1 ppm oxygen concentration.

### Size-exclusion chromatography (SEC) with multi-angle light scattering

SEC was performed using either an S75 Increase 10/300 GL or S200 Increase 10/300 GL column attached to a Dionex HPLC System, PSS SLD7000 Multi Angle Light Scattering Photometer and Shodex RI-101 Control (Data Apex) differential RI detector instruments. The system was calibrated using a 2 mg/mL BSA sample (ThermoFisher). Protein samples (100 μL) were loaded and MW calculations determined using PSS WinGPC UniChrom 8.1 and Thermo Scientific™ Dionex™ Chromeleon™ 7.2 Chromatography Data System software.

### Inductively coupled plasma mass spectrometry

Anaerobically purified SufU samples at 3–5 mg/ml and buffer-matched controls were treated with 2% HNO_3_ and proteins denatured by heating to 70°C in a water bath for 15 minutes. The solutions were quantitatively analyzed on an Agilent 7700 ICP-MS in He-mode to reduce polyatomic interference. Calibration standards were prepared in a matrix matched solution from 1000 ppm Single element standards (CPI International, USA).

### UV/Vis spectrophotometry

A Cary400 spectrophotometer was used to scan SufU and SufR samples containing 1 mg/mL protein in the UV/Vis range (250–600 nm).

### Mössbauer spectroscopy

All samples were prepared under oxygen-free conditions in an anaerobic glovebox (Coylab). Mössbauer spectra were recorded on WEB Research (Edina, MN) instrument. The spectrometer used to acquire the weak-field spectra is equipped with a Janis SVT-400 variable-temperature cryostat. The external magnetic field (if present) was applied parallel to the γ beam. All isomer shifts are quoted relative to the centroid of the spectrum of α-iron metal at room temperature. Mössbauer spectra were fitted using the WMOSS4 Mössbauer Spectral Analysis Software (www.wmoss.org).

### **X-** ray crystallography

Aerobically purified *Mtb*-SufS and zinc-bound *Mtb*-SufU (1:1 molar ratio) were co- crystallized with L-Cys by sitting drop vapor diffusion in a Morpheus crystallization screen^37^ containing 0.1 M carboxylic acids, Morpheus buffer system 1 at pH 6.5, and Morpheus 30% precipitant mix 1. [2Fe-2S]-SufU was crystallized at 7.4 mg/mL in 0.2 M calcium acetate hydrate, 0.1 M sodium cacodylate (pH 6.5), and 40% v/v PEG 3000. Diffraction experiments were carried out at the Australian Synchrotron MX2 beamline. The diffraction image visualization software ADXV^38^ was used to visualize protein diffraction during and after data collection. Data indexing was carried out using X-Ray Detector Software (XDS).^39^ The integrated reflection data was then scaled using AIMLESS^40^ from the CCP4 suite.^41^ 5% of reflections were reserved and maintained for calculation of the R_free_ factor. Molecular replacement was performed using Phenix^42^ and the Alphafold2^43^ generated predicted structure models of *Mtb*-SufS and *Mtb*-SufU. Iterative refinement was done using REFMAC5.^44^ Visual inspection of the crystal packing, electron density, and real-space refinement was carried out using COOT.^45^ Figures and structural alignments were produced using the PyMOL Molecular Graphics System, Version 1.2r3pre, Schrödinger, LLC.

### Cysteine desulfurase assay

The cysteine desulfurase activity of SufS in the presence of zinc- and cluster-bound SufU was determined by analyzing the production of L-Ala. 50 μL reactions consisted of 10 µM SufS, 20 µM SufU, and 1 mM L-Cys in a reaction buffer containing 25 mM HEPES (pH 7.5), 100 mM NaCl, and 5 mM DTT. Reactions were incubated at room temperature under anaerobic conditions, then the reaction was stopped by adding 400 µL 1 M NaOH. Samples were then derivatized by methyl chloroformate alkylation as described in Smart *et al*.^46^ and analyzed by Gas Chromatography-Mass Spectroscopy (GC-MS), utilizing a GC7890A/MSD5975C Gas Chromatograph/mass selective detector system (Agilent Technologies). MassHunter Workstation software (Version 10.0; Agilent Technologies) was used to analyze the results.

### *In vitro* Fe-S cluster assembly

Enzymatic Fe-S reconstitution reactions containing 10 μM *Mtb*-SufS and 50 μM *Mtb*-SufU were pre-equilibrated in a buffer containing 25 mM HEPES (pH 7.5), 100 mM NaCl and 1 mM DTT, before adding FeCl_2_. The reaction was initiated by adding 2 mM L-Cys, and reaction progression was monitored using UV/Vis spectroscopy (Cary400, Varian).

### Treatment of SufU and SufR with H_2_O_2_

Anaerobically purified *Mtb*-SufU and *Mtb*-SufR proteins at 100 μM were incubated with various concentrations of hydrogen peroxide (H_2_O_2_) for two mins before UV/Vis spectrophotometry (Cary400, Varian) was used to assess cluster disruption.

## Acknowledgments

We thank Dr. Jamie Taka, Dr. Matthew Sullivan, and Stuart Morrow for technical assistance. Crystal data collection was undertaken on the MX2 beamline at the Australian Synchrotron. Access to the Australian Synchrotron was supported by the New Zealand Synchrotron Group Ltd.

## Author contributions

Conceptualization: SMS, MP, GB

Methodology: SMS, JC, JW

Investigation: SMS, JC, JW

Funding acquisition: SMS, GB

Project administration: GB

Supervision: SD, ENB, CJS, GB

Writing – original draft: SMS, GB

Writing – review & editing: SMS, JC, JW, SD, ENB, CJS, MP, GB

## Data and materials availability

The coordinates and structure factors for the Zn^2+^-bound *Mtb*-SufS/SufU and [2Fe-2S]-bound *Mtb*-SufU have been deposited in the Protein Data Bank under the accession codes 9DDD and 9DCL, respectively. All other data and material are available from the corresponding author upon request.

## Supplementary Materials

Figs. S1 to S3

Tables S1 and S2

