## Supplementary information (FigS1-S3), Tables S1-S2) for "Feedback regulation of iron-sulfur cluster biogenesis"

**The PDF file includes:**

Figs. S1 to S3

Tables S1 and S2

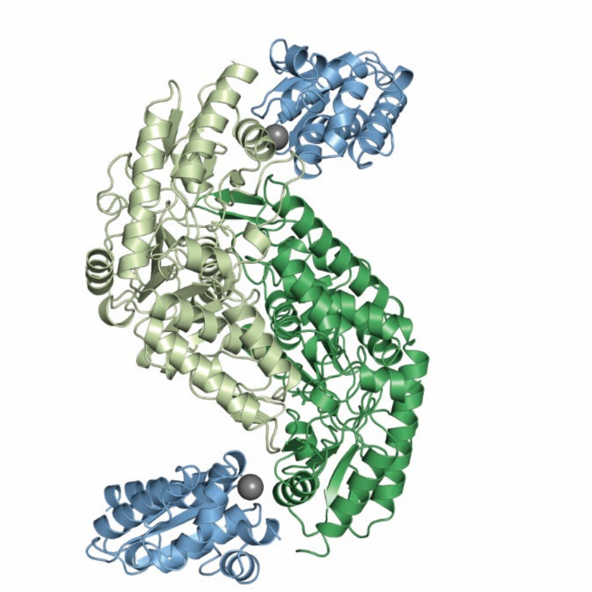

**Fig. S1. Crystal structure of *Mtb*-SufU.** The SufS/SufU complex is a heterotetramer structure, composed of a core SufS dimer (green), with SufU monomers (blue) interacting with each of the SufS chains via a $Zn^{2+}$ ion (gray sphere).

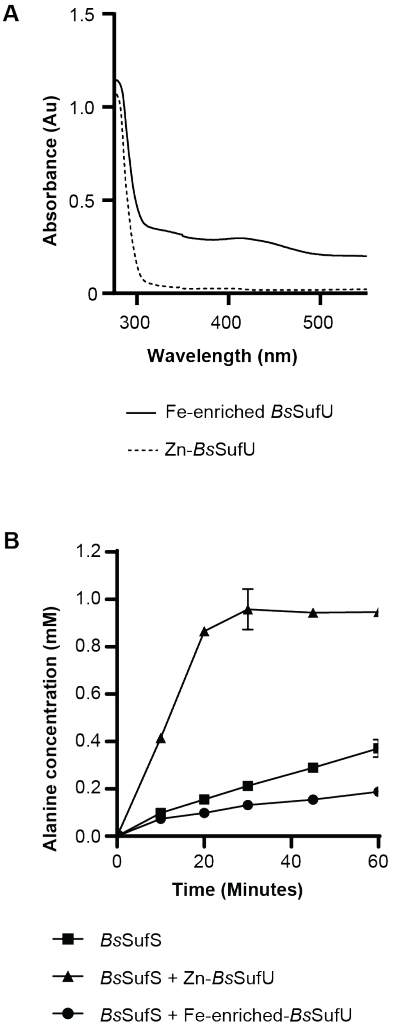

**Fig. S2. Conservation of Fe-S cluster binding in SufU. (A)** UV/Vis absorption spectra of $Zn^{2+}$-bound *Bs*-SufU (dashed line) and as-purified Fe-enriched *Bs*-SufU (solid line). The spectrum of Fe-enriched *Bs*-SufU shows a peak at 420 nm, characteristic of [4Fe-4S] clusters. **(B)** GC-MS cysteine desulfurase activity assay indicates a basal activity of *Bs*-SufS, which is enhanced in the presence of $Zn^{2+}$-bound *Bs*-SufU. While Fe-enriched *Bs*-SufU appears to activate *Bs*-SufS, this is likely due to residual $Zn^{2+}$ (0.2 equivalents per monomer) present in the *Bs*-SufU sample, as confirmed by ICP-MS. Error bars indicate standard deviation in triplicates.

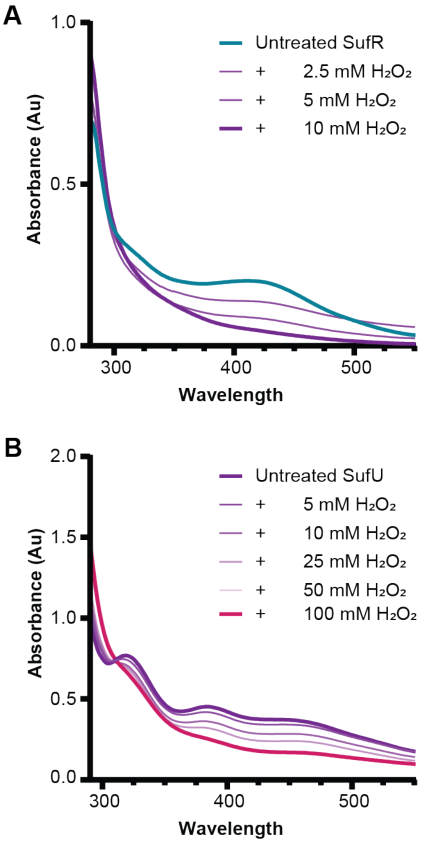

**Fig. S3. SufR and SufU exhibit distinct responses to oxidative stress.** UV/Vis spectra indicates the response of [4Fe-4S]-SufR **(A)** and [2Fe-2S]-SufU **(B)** to various concentrations of $H_2O_2$ two minutes after treatment at each concentration. The [4Fe-4S] cluster in SufR is highly sensitive to oxidation, with its characteristic absorbance signature completely lost after treatment with 10 mM $H_2O_2$. In contrast, the [2Fe-2S] cluster in SufU retains its characteristic features at 50 mM $H_2O_2$.

**Table S1. Summary of ICP-MS analysis of SufU proteins.** The results are presented as equivalent metal ions per protein monomer.

| | **Zinc** | **Iron** |
|---|---|---|
| ***Mtb*-SufU, Zn-bound** | 0.98 | <0.001 |
| ***Mtb*-SufU, [2Fe-2S] bound** | 0.10 | 1.58 |
| ***Sa*-SufU, Fe-enriched** | 1.1 | 0.21 |
| ***Bs*-SufU, Fe-enriched** | 0.24 | 0.48 |

**Table S2. Crystallographic data collection and refinement.**

| | **SufSU** | **SufU** |
|---|---|---|
| **DATA COLLECTION** | | |
| **Source** | Australian Synchrotron | Australian Synchrotron |
| **Wavelength (Å)** | 0.9537 | 0.953725 |
| **Space group** | $P_1$ | $I_4$ |
| **Cell dimensions** | | |
| a, b, c | 66.80, 73.62 ,76.35 | 100.12, 100.12, 31.33 |
| α, β, γ | 91.03, 100.22, 110.02 | 90, 90, 90 |
| **Reflections** | | |
| measured | 645364 (29649) | 294628 (22241) |
| unique | 88203 (4246) | 10785 (796) |
| **Resolution** | 48.65 – 2.00 (2.04 – 2.00) | 35.40 – 2.00 (2.05 – 2.00) |
| **CC½** | 0.998 (0.808) | 0.995 (0.699) |
| **Mean I/σI** | 10.8 (2.5) | 7.8 (1.6) |
| **Completeness** | 98% (91.9%) | 100 (100) |
| **Redundancy** | 7.3 (7.0) | 27.3 (27.9) |
| **Av. Mosaicity** | 0.07 | 0.12 |
| **REFINEMENT** | | |
| **Resolution** | 48.68 - 2.00 | 35.42 - 1.95 |
| **$R_{work}$/$R_{free}$** | 0.180/0.203 | 0.179/0.210 |
| **No. atoms** | | |
| protein | 8320 | 1110 |
| water | 336 | 36 |
| metals/cluster | 2 (Zn) | 4 (2Fe, 2S) |
| PLP | 48 | - |
| Ala | 12 | - |
| ***B*-factors** | | |
| protein | 35.4 | 32.0 |
| water | 35.6 | 34.3 |
| metals/cluster | 35.4 | 22.76 |
| PLP | 25.7 | - |
| Ala | 43.2 | - |
| **R.M.S.D.** | | |
| Bond lengths (Å) | 0.005 | 0.004 |
| Bond Angles (°) | 1.009 | 1.289 |
| **Ramachandran favored** | 98.0% | 100% |
| **Favored rotamers** | 97.3% | 97.4% |
| **Molprobity score** | $100^{th}$ percentile | $100^{th}$ percentile |
